# Mapping migration in a songbird using high-resolution genetic markers

**DOI:** 10.1101/007757

**Authors:** Kristen Ruegg, Eric C. Anderson, Kristina L. Paxton, Vanessa Apkenas, Sirena Lao, Rodney B. Siegel, David F. DeSante, Frank Moore, Thomas B. Smith

## Abstract

Neotropical migratory birds are declining across the Western Hemisphere, but conservation efforts have been hampered by the inability to assess where migrants are most limited – the breeding grounds, migratory stopover sites, or wintering areas. A major challenge has been the lack of an efficient, reliable, and broadly applicable method for connecting populations across the annual cycle. Here we show how high-resolution genetic markers can be used to identify populations of a migratory bird, the Wilson’s warbler (*Cardellina pusilla*), at fine enough spatial scales to facilitate assessing regional drivers of demographic trends. By screening 1626 samples using 96 single nucleotide polymorphisms (SNPs) selected from a large pool of candidates (∼450,000), we identify novel region-specific migratory routes and timetables of migration along the Pacific Flyway. Our results illustrate that high-resolution genetic markers are more reliable, accurate, and amenable to high throughput screening than previously described tracking techniques, making them broadly applicable to large-scale monitoring and conservation of migratory organisms.

## Introduction

Over half of the Neotropical migrant bird species found breeding in North America have shown marked declines in abundance over the last several decades (Robbins 1989; Sauer *et al.* 2012). Population declines are thought to relate to stressors encountered by migrants at each stage in the annual cycle – the breeding grounds, the wintering grounds, and migratory stopover points (Rappole 1995). At each stage birds are subject to a number of disturbances including habitat loss, collisions with wind turbines and cell phone towers, predation by house cats, exposure to disease, and the increasing effects of global climate change (Altizer *et al.* 2011; Jonzen *et al.* 2006; Loss *et al.* 2013). However, without the ability to connect populations across the annual cycle it is difficult to assess the impact of local stressors on population declines. Historically, efforts to map songbird migration patterns relied on recovery of individual birds previously captured and tagged with bird bands; however, this approach has met with limited success for small-bodied songbirds because recapture rates of birds away from their original banding sites are often very low (< 1 in 10,000) (Faaborg *et al.* 2010b; Gustafson & Hildenbrand 1999). More recently, geo-locators, small tracking devices that record information on ambient light levels to estimate an individuals location, have increased our knowledge of the migratory pathways in many songbird species (Stutchbury *et al.* 2009), but remain impractical for most large-scale applications (1000’s of individuals) due to cost, weight restrictions, and the need to recover individuals to collect data from the devices (Arlt *et al.* 2013; Bridge *et al.* 2013). Alternatively, genetic and isotopic markers that use information contained within the feathers to pinpoint an individuals population of origin have broad appeal because they are cost-effective, noninvasive, and do not require recapture (Kelly *et al.* 2005; Rubenstein *et al.* 2002; Rundel *et al.* 2013), but have been plagued in the past by low resolution and/or technical issues related to working with feathers (Lovette *et al.* 2004; Segelbacher 2002; Wunder *et al.* 2005). Thus, there remains a need for a broadly applicable tracking method that can be used to resolve populations on spatial scales that are informative for assessing drivers of regional population declines.

In the last several years, genome sequencing has revolutionized the field of molecular ecology, resulting in new technologies that can be applied to molecular tagging of wild populations (Davey *et al.* 2011). Genome reduction techniques, such as Restriction Site Associated DNA sequencing (RAD-seq), can be used to sequence multiple individuals across a large fraction of the genome and identify hundreds of thousands of genetic markers that are useful for distinguishing populations (Baird *et al.* 2008). One type of genetic marker that can be identified from genomic sequence data is a Single Nucleotide Polymorphism (SNP), DNA sequence variation occurring when a single nucleotide in the genetic code – A, T, C, or G – differs between individuals or homologous chromosomes. In particular, SNPs found within or linked to genes under selection often display elevated allele frequencies and, as a result, can be targeted to reveal population structure at finer spatial scales than is possible using neutral genetic markers (Nielsen *et al.* 2012; Nielsen *et al.* 2009). Furthermore, SNP-specific assays designed to target small fragments of sequence around the SNP loci of interest can be advantageous in cases where the DNA is highly fragmented or available only in very small quantities, such as DNA from a single, small passerine feather.

Here we develop high-resolution SNP markers for tracking populations of a migratory bird, the Wilson’s warbler, *Cardellina pusilla,* using a combination of Restriction Site Associated DNA paired end sequencing (RAD-PE seq) and high throughput SNPtype^TM^ Assay screening. The Wilson’s warbler, a long-distance neotropical migratory bird with a cross-continental breeding distribution (Ammon & Gilbert 1999), is particularly appropriate as model for testing the efficacy of high-resolution molecular markers because previous population genetic/connectivity studies on this species provide a solid basis for comparison between methods (Clegg *et al.* 2003; Irwin *et al.* 2011; Kimura *et al.* 2002; Paxton *et al.* 2007; Paxton *et al.* 2013; Rundel *et al.* 2013; Yong *et al.* 1998). By harnessing recent advances in Next-Generation Sequencing we scan the genomes of Wilson’s warblers sampled from across the breeding range and identify a set of highly divergent SNP loci with strong potential for population identification. We then develop SNPtype^TM^ Assays that target these highly divergent loci and use them to screen 1626 feather and blood samples collected from across the annual cycle in collaboration with bird banding stations located across North and Central America. We illustrate how the resulting region-specific migration map can be used to help identify drivers of regional demographic trends and inform studies of migrant stopover ecology.

## Methods

### (a) Sample collection

Collection of 1648 feather and blood samples (22 samples for the SNP ascertainment panel and 1626 for the SNP screening panel) from 68 locations across the breeding, wintering and migratory range was made possible through a large collaborative effort with bird banding stations within and outside of the Monitoring Avian Productivity and Survivorship (MAPS), the Landbird Monitoring of North America (LaMNA), and the Monitoreo de Sobrevivencia Invernal (MoSI) networks (Table 1). Genetic samples, consisting of the tip of one outer rectrix or blood collected by brachial vein puncture and preserved in lysis buffer (Seutin 1991), were purified using Qiagen DNeasy Blood and Tissue Kit and quantified using a NanoDrop™ Spectrophotometer (Thermo Scientific, Inc) (Smith *et al.* 2003). Breeding (June 10 – July 31), migratory (March 1 – May 31), and wintering (December 1 – February 28) samples were collected and categorized into groups based on collection date, signs of breeding (presence/size of a cloacal protuberance), signs migration (extent of fat) and life history timetables for the Wilson’s warbler (Ammon & Gilbert 1999). To assess migratory stopover site use through time, 686 of the 1648 samples from a stopover site located on the Lower Lower Colorado River, near the town Cibola, AZ, were collected using consistent effort, daily, passive mist-netting from March 22 – May 24, across the years 2008 and 2009 (Table 1).

**Table 1.**
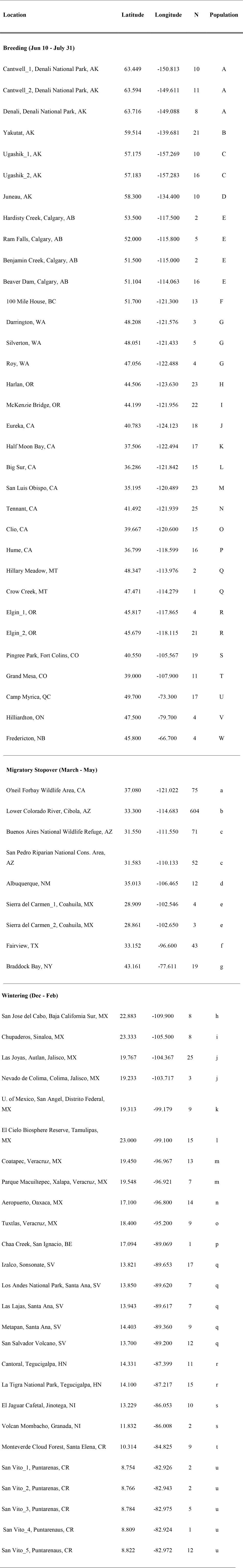
Number of Wilson’s warblers successfully screened at each location across the species breeding, wintering and migratory range. Locations in close proximity were merged on the map in Fig. 1. Uppercase letters are reserved for breeding populations, while lower case letters are reserved for migratory stopover and wintering locations.

### (b) SNP discovery

To identify SNPs useful for distinguishing genetically distinct regions across the breeding range of the Wilson’s warbler, an ascertainment panel of 22 individuals was selected to represent the range of phylogenetic variation known in the species, including all 3 recognized subspecies (Ammon & Gilbert 1999; Kimura *et al.* 2002). Five individuals from each of five regions were included in the ascertainment panel, except for from the Southwestern region where samples were limited to 2 individuals (SI Table 1). Purified extractions from blood samples were quantified using Quant-iT™ PicoGreen^®^ dsDNA Assay Kit (Invitrogen Inc), and Restriction Site Associated DNA Paired-end (RAD-PE) libraries containing individually barcoded samples were prepared at Floragenex, Inc. according to Baird *et al.* (2008) and Ruegg *et. al*. (Ruegg *et al.* 2014) (SI Methods). RAD-PE sequencing made it possible to build longer contigs (∼300bp) from short read, 100bp Illumina HiSeq2000 data in order to improve downstream bioinformatics and provide adequate flanking sequence around SNPs for assay development (Etter *et al.* 2011).

Samples from each isolate were sequenced on an Illumina HiSeq2000 (Illumina, San Diego, CA) using paired-end 100 bp sequencing reads. Paired-end sequences from each sample were collected, separated by individual, stripped of barcodes, trimmed to 70 bp, scrubbed of putative contaminant and high-copy-number-sequences and filtered to include only those with a Phred score ≥10. The sample with the greatest number of reads passing the initial quality filter was used to create a reference set of RAD-PE contigs against which sequences from other samples were aligned. To create the reference, primary reads were clustered into unique RAD markers and the paired-end sequences associated with each RAD tag were assembled *de novo* using Velvet (Zerbino & Birney 2008) into contigs ranging from 180 – 610 bp, with an average length of 300bp. Paired-end reads from the remaining samples were aligned to this reference using Bowtie (Langmead & Salzberg 2012) and SNPs were identified using the SAMtools software (Li *et al.* 2009) with mpileup module under standard conditions.

To narrow our dataset to SNPs we could confidently use to assess population structure we performed a second round of quality filtering and removed: (1) putative SNPs with no variants and / or more than two alleles; (2) genotypes in individuals with a quality score of < 30; (3) genotypes with < 8 reads in a homozygote or < 4 reads per allele in heterozygotes; (4) putative SNPs that had suitable genotypes in < 12 out of the 17 samples from four western populations or < 5 out of the 5 samples from the eastern population and (5) putative SNPs with < 40 bp of flanking sequence on either side. To limit the chances of including linked markers genomic coordinates were attained by mapping the remaining contigs to the closest, best annotated, songbird genome, the zebra finch (*Taeniopygia guttata*) (Version 3.2.4; (Warren *et al.* 2010)) using BLAST+ (version 2.2.25).

To avoid the possibility of erroneous matches, the data was filtered to include only contigs that aligned to the zebra finch genome with only a single hit and an E-value < 10^−40^. Because SNPs with large frequency differences are the most effective for identifying populations, all SNPs that passed our second round of quality filters were ranked according to frequency differences between the 5 regions (SI Table 2) and 150 SNPs displaying the largest allele frequency differences between each of the 10 pairwise comparisons were selected for conversion to SNPtype^TM^ Assays (Fluidigm Inc). Before making a final selection, we also considered factors such as: GC content (<65%), number of genotypes per population, and average coverage at a SNP across all populations (SI Table 2). An initial assay pre-screening panel was then performed and the assay pool was further reduced to the 96 assays (the number that fit on a single 96.96 Fluidigm Array) that could be genotyped most reliably (SI Table 2).

**Table 2.** Assignment of Wilson’s warblers of known origin back to breeding population using GSI_Sim. Population names are listed in Table 1 and the colors indicate the genetic group (Fig. 1).

| **Population**<br>**(Fig. 1,**<br>**Table 1)** | **Alaska to**<br>**Alberta** | **Pacific**<br>**Northwest** | **Coastal**<br>**California** | **Sierra** | **Rocky**<br>**Mountain** | **Eastern** |
| --- | --- | --- | --- | --- | --- | --- |
| **A** | 29 | 0 | 0 | 0 | 0 | 0 |
| **B** | 21 | 0 | 0 | 0 | 0 | 0 |
| **C** | 26 | 0 | 0 | 0 | 0 | 0 |
| **D** | 10 | 0 | 0 | 0 | 0 | 0 |
| **E** | 24 | 0 | 0 | 0 | 1 | 0 |
| **F** | 9 | 0 | 0 | 0 | 4 | 0 |
| **G** | 2 | 9 | 1 | 0 | 0 | 0 |
| **H** | 0 | 20 | 3 | 0 | 0 | 0 |
| **I** | 0 | 20 | 1 | 1 | 0 | 0 |
| **J** | 0 | 15 | 0 | 3 | 0 | 0 |
| **K** | 0 | 2 | 14 | 1 | 0 | 0 |
| **L** | 0 | 3 | 11 | 1 | 0 | 0 |
| **M** | 0 | 1 | 19 | 3 | 0 | 0 |
| **N** | 0 | 2 | 2 | 21 | 0 | 0 |
| **O** | 0 | 1 | 2 | 12 | 0 | 0 |
| **P** | 0 | 1 | 0 | 15 | 0 | 0 |
| **Q** | 1 | 0 | 0 | 0 | 2 | 0 |
| **R** | 6 | 0 | 0 | 0 | 19 | 0 |
| **S** | 0 | 0 | 0 | 0 | 19 | 0 |
| **T** | 0 | 0 | 0 | 0 | 11 | 0 |
| **U** | 0 | 0 | 0 | 0 | 0 | 17 |
| **V** | 0 | 0 | 0 | 0 | 0 | 4 |
| **W** | 0 | 0 | 0 | 0 | 0 | 4 |

### (b) SNP Screening

The Fluidigm Corporation EP1™ Genotyping System was used to genotype 96 SNP loci using 94 individuals per run and 2 non-template controls. To avoid the potential for high-grading bias (i.e. wrongly inflating the apparent resolving power of a group of loci for population identification) (Anderson 2010), none of the 22 samples used in our original ascertainment panel were included in the final SNP screening and population structure analyses. To ensure amplification of low quality or low concentration DNA from feathers, an initial pre-amplification step was performed according to the manufacturers protocol using a primer pool containing 96 unlabeled locus-specific SNPtype primers (SI Methods). PCR products were diluted 1:100 and re-amplified using fluorescently labeled allele-specific primers. The results were imaged on an EP1 Array Reader and alleles were called using Fluidigm’s automated Genotyping Analysis Software (Fluidigm Inc) with a confidence threshold of 90%. In addition, all SNP calls were visually inspected and any calls that did not fall clearly into one of three clusters – heterozygote or either homozygote cluster - were removed from the analysis. As DNA quality can affect call accuracy, a stringent quality filter was employed and samples with >90 of 96 missing loci were dropped. To assess the reliability of SNPtype assays for genotyping DNA from a variety of sources (blood and feather extractions), the proportion of samples yielding useable genotype data was calculated. Tests for linkage disequilibrium and conformance to Hardy-Weinberg equilibrium (HWE) (Louis & Dempster 1987) were performed using GENEPOP software, vers. 4.0 (Rousset 2008).

### (c) Population structure analysis

While genetic differentiation (F_ST_) is likely inflated because selected loci were not a random sample from the genome, we calculated F_ST_ here for comparison to previous genetic analysis. F_ST_ between all pairs of populations was calculated as θ (Weir & Cockerham 1984), using the software GENETIX vers. 4.05 (Belkhir *et al.* 1996-2004) and the data were permuted 1000 times to determine significance. We used the program STRUCTURE ver. 2.2, to further assess the potential for population structure across the breeding grounds (Pritchard *et al.* 2000). Ten runs at each K value (K= 1-9) were performed under the admixture model with correlated allele frequencies using a burn-in period of 50,000 iterations, a run length of 150,000. All scripts used for the STRUCTURE runs and subsequent population genomic analyses are located at https://github.com/eriqande/wiwa-popgen. To simplify comparison of results, the program CLUMPP (Jakobsson & Rosenberg 2007) was used to reorder the cluster labels between runs, and individual *q* values (proportion of ancestry inferred from each population within an individual) were plotted using the program Distruct (Rosenberg 2004). Visual inspection of Distruct plots allowed identification of regions where geographic barriers to gene flow exist and/or where admixture is likely.

To identify how population structure was distributed across geographic space, we used the program GENELAND (Guillot *et al.* 2005). Analyses in GENELAND were performed under the spatial model assuming uncorrelated allele frequencies. Inference of population structuring was based on 10 independent runs, each allowing the number of populations to vary between 1 and 10. Each run consisted of 2.2 million MCMC iterations with a thinning interval of 100. Of the 22,000 iterations retained for the MCMC sample after thinning, the first 5,000 were discarded as burn-in. Post processing of the MCMC sample was done upon a 250 by 250 point grid that covered the breeding range of the species. Posterior probability of group membership estimates from GENELAND were visualized as transparency levels of different colors overlaid upon a base map from Natural Earth (naturalearthdata.com) and clipped to the Wilson’s warbler breeding range using a shapefile (NatureServe 2012), making use of the packages sp, rgdal, and raster in R (Bivand *et al.* 2014; Hijmans 2014; Pebesma & Bivand 2005; Team 2014) (see https://github.com/eriqande/wiwa-popgen). Thus, within each distinguishable group the transparency of colors is scaled so that the highest posterior probability of membership in the group according to GENELAND is opaque and the smallest is entirely transparent.

To assess the accuracy of our baseline for identification of individuals from each population to genetically distinct breeding groups we used the program GSI_Sim (Anderson 2010; Anderson *et al.* 2008). GSI_Sim uses an unbiased leave-one-out cross-validation method to assess the accuracy of self-assignment of individuals to populations. Posterior probabilities were obtained in GSI_Sim by summing the posterior probabilities of the populations within each genetically distinct group and assigning the individual to the genetically distinct group with the highest posterior probability.

## Results

### (a) SNP discovery

RAD-PE sequencing on 22 individuals from 5 geographic regions representative of the range of phylogenetic variation known in the species resulted in 123,005 contigs (average length ∼300 bp), containing 449,596 SNPs passing our initial quality filters (SI Table 1). The median depth of sequencing across all contigs within a library was 33x and the average Phred quality score per library was 35 (SI Table 1). Overall, 166,268 SNPs passed the second round of quality filters and 19,707 of those were candidates for conversion into SNPtype™ Assays based upon the absence of variation in 40 base pairs of flanking sequence surrounding the SNPs. Candidate SNPs were ranked according to frequency differences, GC content, the number of genotypes per region, and the average coverage and the final panel was composed of 96 SNPs with pairwise frequency differences between regions ranging from 1 – 0.4 (SI Table 2). For contigs that could be mapped to the zebra finch genome with high confidence, the minimum distance between SNPs was 41KB and no two SNPs were selected from the same contig in order to avoid the possibility of linked markers (SI Table 2). In this study we refer to the final panel of 96 highly differentiated SNPs as high-resolution genetic markers.

### (b) SNP screening

The resulting high resolution genetic markers were used to screen 1626 samples collected from 68 sampling locations across the breeding, wintering and migratory range (Table 1), with 117 samples excluded due to low quality genotypes (>6 loci excluded). The samples with the highest proportion of reliable genotypes were from fresh feather extractions (n_reliable_/ total = 660/686 or 96% reliable), followed by fresh blood extractions (n_reliable_ / total = 100/106 or 94% reliable), and finally extractions that were >3 years old (n_reliable_ / total = 701/786 or 90% reliable). Tests for conformity to HWE revealed that all but 1 of the 94 loci (AB_AK_20) in 2 of the 23 breeding populations (D an L; Table 1, Fig. 1b) were in HWE after accounting for multiple comparisons (p<0.0005). Deviations from HWE were likely the result of small sample sizes and or the unintentional inclusion of late arriving migrants *en route* to northern breeding sites. No loci were found to be in linkage disequilibrium after accounting for multiple comparisons (p<0.0005), suggesting that loci were not physically linked even in cases where zebra finch genome coordinates could not be attained.

**Figure 1.**
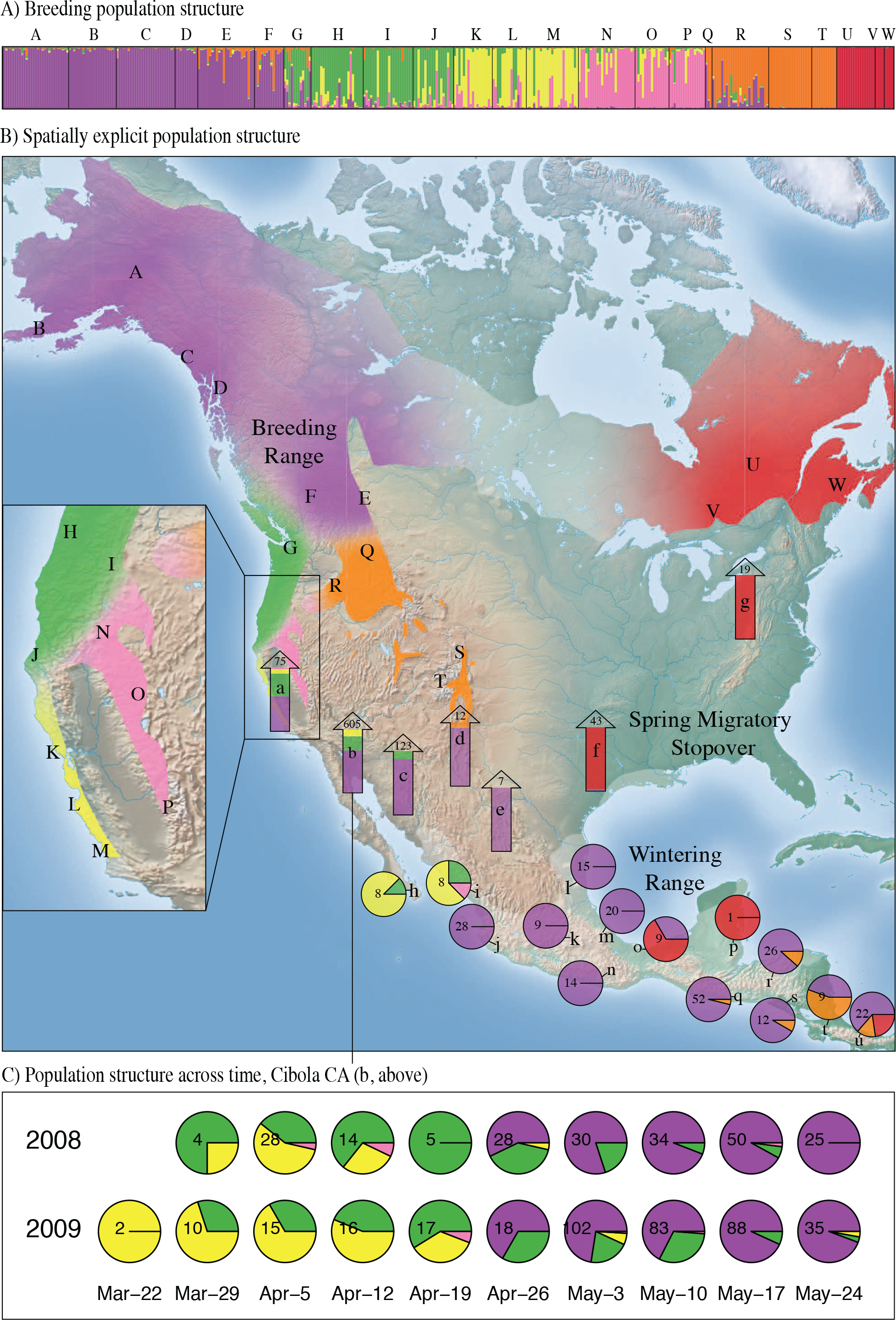
Migratory connections in the Wilson’s warbler identified using SNP-based genetic markers. A) Results from STRUCTURE showing 6 genetically distinct populations across the breeding grounds. Capital letters (A-W) refer to the location of breeding populations depicted on the map in B as well as listed in Table 1. B) Spatially explicit population structure across the annual cycle. The colors across the breeding range represent the results from GENELAND which were post-processed using R so that the density of each color relects the relative posterior probability of membership for each pixel to the most probable of the 6 different genetic clusters (see text). The results were clipped to the species distribution map (NatureServe 2012). Lower case letters (a-g) represent the location of wintering and spring migratory samples (Table 1). Pie charts indicate the proportion of wintering indidiviuals assigned to each breeding group with the number of individuals listed at the center of each pie. Arrows represent the proportion of migrants assigned to each breeding group with the numbers of indviduals listed at the top of the arrows. C) The proportion of indivdiuals assigned to each breeding population across spring migration of 2008 and 2009. Numbers in the center of the pies refer to sample sizes and the data are grouped by week with the date representing the mid-week date in a non-leap year.

### (c) Population Structure analysis

An analysis of population genetic structure on the breeding grounds identified 6 genetically distinguishable groups: Alaska (purple, A - D), eastern North America (red, U-W), the Southern Rockies and Colorado Plateau (orange, S, T), the Pacific Northwest (green, G-J), Sierra Nevada (pink, N-P), and Coastal California (yellow, K-M) (Fig. 1a&b). Pairwise F_ST_’s between groups ranged from 0-0.68 with an overall F_ST_ of 0.179 (95% CI: 0.144 – 0.218). The strongest genetic differentiation was observed between eastern and western groups (F_ST_ = 0.41 – 0.68) with strong genetic differentiation also seen between the Southern Rockies and Colorado Plateau and all other groups (F_ST_ = 0.09 – 0.27; SI Table 3). The number of genetically distinct groups was set at 6 based upon convergence between results from STRUCTURE (k=6, average ln P(X|K) = −33359), GENELAND, and GSI_Sim (Fig. 1a&b; Table 2). While 7 genetically distinct groups was also strongly supported by GENELAND and STRUCTURE (K=7, average ln P(X|K) = −33286; SI Fig. 1), with sampling locations from British Columbia and Alberta (E and F) forming a seventh group distinct from Alaska, the power to accurately assign individuals to groups at k=7 decreased significantly using both STRUCTURE and GSI_Sim (SI Fig. 1).

**Table 3.** Genetic identification of Wilson’s Warblers migrating through Cibola, CA by week across the years 2008 and 2009. Results represent the individuals assigned to one of the six genetically distinct groups using the program GSI Sim and the data corresponds to the information presented in Figure 1c.

| Mid-week Date* | Week | Alaska to Alberta | Pacific NW | Coastal CA | Sierra Nevada | Rocky Mt. | Eastern |
| --- | --- | --- | --- | --- | --- | --- | --- |
| **Year 2008** |  |  |  |  |  |  |  |
| 21-Mar | 11 | 0 | 0 | 0 | 0 | 0 | 0 |
| 28-Mar | 12 | 0 | 3 | 1 | 0 | 0 | 0 |
| 4-Apr | 13 | 0 | 11 | 16 | 1 | 0 | 0 |
| 11-Apr | 14 | 0 | 9 | 4 | 1 | 0 | 0 |
| 18-Apr | 15 | 0 | 5 | 0 | 0 | 0 | 0 |
| 25-Apr | 16 | 16 | 11 | 1 | 0 | 0 | 0 |
| 2-May | 17 | 24 | 6 | 0 | 0 | 0 | 0 |
| 9-May | 18 | 32 | 2 | 0 | 0 | 0 | 0 |
| 16-May | 19 | 46 | 3 | 0 | 1 | 0 | 0 |
| 23-May | 20 | 25 | 0 | 0 | 0 | 0 | 0 |
| **Year 2009** |  |  |  |  |  |  |  |
| 22-Mar | 11 | 0 | 0 | 2 | 0 | 0 | 0 |
| 29-Mar | 12 | 0 | 3 | 7 | 0 | 0 | 0 |
| 5-Apr | 13 | 0 | 5 | 10 | 0 | 0 | 0 |
| 12-Apr | 14 | 0 | 6 | 10 | 0 | 0 | 0 |
| 19-Apr | 15 | 0 | 10 | 6 | 1 | 0 | 0 |
| 26-Apr | 16 | 12 | 6 | 0 | 0 | 0 | 0 |
| 3-May | 17 | 74 | 21 | 6 | 1 | 0 | 0 |
| 10-May | 18 | 56 | 25 | 1 | 1 | 0 | 0 |
| 17-May | 19 | 82 | 6 | 0 | 0 | 0 | 0 |
| 24-May | 20 | 33 | 1 | 1 | 0 | 0 | 0 |

\* Dates represent the midweek date in a non-leap year

Leave-one-out cross validation using GSI_Sim indicated that the ability to correctly assign individuals to groups was high, ranging from 80 - 100%. The eastern group had the highest probability of correct assignment (100%), followed by Alaska to Alberta (94%), the Southern Rockies and Colorado Plateau (92%), the Pacific Northwest (84%), the Sierra Nevada (81%) and Coastal California (80%) (Fig. 1b; Table 2). The majority of the incorrect assignments were between the Pacific Northwest, Sierra Nevada and Coastal California. Subsequent assignment of migrant and wintering individuals to genetically distinct breeding groups using GSI_sim indicated that Coastal California, Sierra, and Pacific Northwest breeders winter in western Mexico and southern Baja, and migrate north along the Pacific Flyway, with Coastal California and Sierra breeders found to the west of the Lower Colorado River (Fig. 1b; SI Table 4). In contrast, Southern Rocky and Colorado Plateau breeders winter from El Salvador to Costa Rica, and migrate north through the central US, while eastern breeders winter primarily in the Yucatan and southern Costa Rica and migrate north through eastern Texas and New York (Fig. 1b; SI Table 4). Unlike the presence of strong connectivity across much of the range, Wilson’s warblers breeding from Alaska to Alberta were identified in all but one of our migratory stopover sites and across all wintering areas, apart from western Mexico and southern Baja (Fig. 1b, all but location g; SI Table 4).

Assignment of migrants collected in a time series from Cibola, AZ revealed a strong temporal pattern in stopover site use across the spring migratory period (Fig. 1c; Table 3). Birds *en route* to coastal California arrived first (week of March 22), followed by birds *en route* to the Pacific Northwest (week of March 29), the Sierra Nevada (week of Apri l 5), and Alaska to Alberta (week of April 26). Only a few individuals migrating through the stopover site were identified as Sierra Nevada breeders (3 per year), while no populations breeding in the Southern Rocky and Colorado Plateau and Eastern U.S. were identified migrating through the stopover site. Temporal patterns in the arrival of spring migrants were replicated across both the years 2008 and 2009 and were consistent regardless of known differences in migration patterns by age and sex (Yong *et al.* 1998).

## Discussion

Full life cycle conservation of declining migrant songbirds has been hindered by lack of an efficient tracking technology that is both broadly applicable and high resolution. Here we demonstrate how high-resolution molecular markers can be applied towards full life cycle conservation of a migrant songbird, the Wilson’s warbler, with a degree of reliability and efficiency that has not been demonstrated using previous tracking methods. By harnessing recent advances in Next-Generation Sequencing we show that 96 highly divergent SNPs selected from a large pool of candidates (∼450,000 SNPs) can be used to identify genetically distinct groups on spatial scales that are informative for regional conservation planning. Our analysis indicates that the power to identify individuals to breeding populations is high (80 - 100%) and that reliable genotypes can be attained from 96% of feathers collected non-invasively from established bird monitoring stations across North and Central America. Because of the biallelic nature of the SNPs in our panel, our genetic data are also easier to validate and standardize across labs than isotope and other genetic methods and, once the assays have been developed, it is possible to genotype ∼300 birds per day for < $10.00/ individual in almost any well-equipped molecular laboratory. Overall, the resolution, efficiency, and cost, combined with the ease of feather collection in collaboration with existing bird monitoring/banding infrastructure, makes high-resolution genetic markers a broadly applicable method for widespread monitoring of declining songbird species.

One of the central challenges in migratory bird conservation is that population declines and conservation planning often occur at regional spatial scales, but our knowledge of migratory connections is usually limited to species-wide range maps. For example, in the Wilson’s warbler, an analysis of Breeding Bird Survey (BBS) data for the years 1966 – 2012 suggests that the species is only slightly declining across it’s range (BBS Trend = - 1.88, 95% CI = 2.97, −1.11), but an analysis of regional trends suggest that populations in the Sierra Nevada and the Southern Rockies/Colorado Plateau are declining more strongly (BBS Trend_sierra_ = −4.71, 95% CI = −6.41, −2.85; BBS Trend_rockies_ = −2.95, 95% CI = −4.32, −1.42) (Sauer *et al.* 2012). Here we illustrate that by targeting highly divergent SNP loci we can confidently identify a minimum of six genetically distinct groups across the breeding range with a resolution in the western US equivalent to the spatial scale of regional population declines. Furthermore, the spatial scale of our genetic groups is commensurate with many *a priori* defined Bird Conservation Regions, ecologically distinct areas in North America with similar habitats and resource management issues (Millard *et al.* 2012). The ability to align the spatial scale of population genetic structure with the spatial scale of population declines and conservation planning provides a powerful framework from which to base full life cycle conservation (Fig. 1a & b).

The Wilson’s warbler has been the focus of numerous population genetic/connectivity studies in the past decade (Clegg *et al.* 2003; Irwin *et al.* 2011; Kimura *et al.* 2002; Paxton *et al.* 2007; Paxton *et al.* 2013; Rundel *et al.* 2013; Yong *et al.* 1998), but none have yielded the depth and clarity of information on migratory connections documented herein. Our results confirm the presence of previously identified connections between birds breeding in Coastal California and wintering in Southern Baja, MX and between birds breeding in eastern North America and wintering in the Yucatan, Belize and Costa Rica (Kimura *et al.* 2002; Rundel *et al.* 2013), but also reveal new patterns across time and space that are much richer and stronger then previously recognized. For example, here we show that Wilson’s warblers breeding in Coastal California (Fig. 1b, yellow) share their wintering area in southern Baja with Pacific Northwest breeders (Fig. 1b, green) and that both of these groups also winter to the east of Baja in Sinaloa, MX, with Sierra Nevada breeders (Fig. 1b, pink) (Sauer *et al.* 2012). Samples collected from across the spring migratory period indicate that western breeders from all three groups (Coastal California, Pacific Northwest, and Sierra Nevada) migrate north along the Pacific Flyway, with Coastal California and Sierra Nevada breeders found west of the Lower Colorado River. In addition, we show for the first time that breeders from the Southern Rocky Mountains and Colorado Plateau (Fig. 1b, orange) occupy a restricted El Salvador-to-Costa Rica wintering distribution and migrate North along the Central Flyway, while eastern breeders (Fig. 1b, red) migrate North through eastern Texas and New York. Overall our results indicate that screening high volumes of individuals using high resolution molecular markers can yield a level of clarity in migratory connections across time and space that has not been previously demonstrated using other tracking techniques.

The resulting map for the Wilson’s warbler provides an example of how information on region-specific migration patterns can be combined with information on region-specific population declines in order to strengthen predictions about where migrants are most limited. In the case of the Wilson’s warbler, BBS data suggests that Sierra Nevada breeders are experiencing strong population declines (BBS Trend_sierra_ = 4.71, 95% CI = - 6.41, −2.85), while Pacific Northwest and Coastal California breeders are declining less severely or remaining stable (BBS Trend_Pacific_Northwest_ = −1.96, 95% CI = −2.54, −1.31; BBS Trend_Coastal_California_ = −0.49, CI = −1.62, 0.84). The fact that all three groups occupy distinct breeding ranges, but mix on their wintering grounds and at migratory stopover sites suggests that declines in Sierra Nevada breeders are likely driven by factors on the breeding grounds. Alternatively, the migration map as a whole suggests that bottlenecks for Wilson’s warblers likely occur in areas where multiple genetically distinct breeding groups funnel through the same stopover site or wintering area such as in Coastal California, Western Mexico, and Costa Rica. These results are supported by work in other taxa and further emphasize the importance of stopover habitat for migrant conservation (Sheehy *et al.* 2011).

Migratory passerines spend roughly a quarter of their year *en route* between breeding and wintering areas, but relatively little is known about the biology and behavior of migrants during the migratory phase of their annual cycle (Faaborg *et al.* 2010b). The availability and quality of habitat at stopover sites could have significant effects on populations, but determining the extent to which physiological and ecological demands experienced during migration may limit populations is often contingent upon knowledge of an individuals ultimate destination (Faaborg *et al.* 2010a; Faaborg *et al.* 2010b). Here we successfully genotype 609 samples collected in a time series from a stopover site near Cibola, AZ and demonstrate how high-resolution genetic markers can be used to identify the ultimate destination of birds captured *en route* to their breeding grounds (Fig. 1b &c; location b). Breaking down the results by week revealed distinct waves of migrants, with Coastal California breeders arriving first (March 22 – 29), followed by Pacific Northwest and Sierra Nevada breeders (March 29-April 5), and Alaska-to-Alberta breeders arriving significantly later (April 19-26). These patterns were replicated across two years and are consistent regardless of known differences in migration patterns by age and sex (Yong *et al.* 1998). While differences in the timing of migration in Wilson’s warblers have been suggested in the past based upon changes in the frequency of haplotypes or isotopic signatures (Paxton *et al.* 2007; Paxton *et al.* 2013), this is the first time that anyone has attained individual-level assignments of large numbers of migrants collected in a time series, bringing a new level of clarity to our understanding of stopover site use through time. It is important to note, that the depth of sampling across time that we are able to achieve using high-resolution genetic markers would not have been possible using extrinsic tracking devices, such as geolocators, due of cost and weight restrictions and the need to recapture individuals to collect the information (Arlt *et al.* 2013; Bridge *et al.* 2013). The resulting information on migratory connections across time can be used to help build timetables of migration along the Pacific Flyway and help to inform when particularly vulnerable populations may be migrating through an area. Furthermore, because DNA can be collected from all birds, dead or alive, high resolution genetic markers could be used to identify migrants subject to collisions with wind turbines, cell phone towers and other manmade structures.

While our results suggest that high-resolution molecular markers surpass previous genetic markers in terms efficiency and resolution, our conclusions could be further strengthened by the inclusion of additional data and analyses. For example, the robustness of the patterns described here varies depending upon the sample size at each location and in some locations, such as in Belize and many of the migratory stopover sites (Fig. 1b, locations l, d, e, f, g), additional sampling across time and space is needed. In addition, while our assignment probabilities are very high for an intrinsic marker (80 – 100%) there is a potential for incorrect assignments, particularly between the three western groups (Coastal California, Pacific Northwest, and the Sierras) were admixture is likely (Table 2). Similarly, there are large regions on the breeding grounds that could not be distinguished using our markers, such as birds breeding from Alberta to Alaska (purple, Fig. 1b). In the future, the addition of more genetic loci as well as the addition of isotopic markers and statistical methods for combining both sources of data into a single statistical framework will help further resolve populations across the range (Rundel *et al.* 2013). Lastly, it is important to note that the spatially explicit depiction of the genetic results generated in GENELAND may not accurately identify the location of boundary between genetic groups. Additional sampling across the projected boundaries will help clarify the location of the genetic breaks as well as the factors driving differences between Wilson’s warblers in each region. Such genetic differences are particularly interesting in light of the documented differences in migratory timing for Wilson’s warblers described herein and the potential for migration timing to contribute to divergence in migratory birds more generally (Bearhop *et al.* 2005; Ruegg *et al.* 2014; Ruegg *et al.* 2012).

A review article by Faaborg *et al* (Faaborg *et al.* 2010b) recently identified continuing research needs for Neotropical migrant birds, including identifying migratory pathways and wintering locations, bottlenecks for conservation, and timetables for migration. Here we demonstrate how high-resolution genetic markers designed for Wilson’s warblers, can be applied to help address many of these continuing research needs with a level of efficiency and reliability that has not previously been demonstrated. In the last several years there has been a revolution in sequencing technology that has increased by orders of magnitude the amount of sequence data that can be generated, while at the same time reducing the cost of individual-level analysis (Metzker 2010). Our results show that by harnessing recent advances in sequencing technology it is now possible to develop high-resolution genetic markers for tracking populations of migrants on a broad scale. The resulting information on fine-scale population genetic structure, region-specific migratory connections, and timetables of migration provides a powerful framework from which to base full life cycle conservation of declining songbird species.

## Acknowledgements

We would like to thank J.C. Garza at the Southwest Fisheries Science Center for the use of laboratory space and equipment. This research was supported by a grant to K. Ruegg from the California Institute for the Energy and the Environment (POEA01-Z01), a donation from Margery Nicolson, a grant to T.B. Smith from The Turner Foundation, EPA (RD-83377801), and a grant to F. Moore from the National Science Foundation (IOS-0844703). We would also like to thank J. Boone and T. Atwood for their assistance with the laboratory and bioinformatics components of the assay design, the many LaMNA, MAPS, and MoSI station operators who contributed avian tissue samples, C.J. Ralph, L. West, D. Kaschube, P. Pyle, J. Saracco, and R. Taylor for coordinating sampling efforts. We are grateful to the USDI Fish and Wildlife Service, the National Park Service, and USDA Forest Service for funding to help operate LaMNA, MAPS and MoSI stations that provided feather samples for this work as well as field crews and staff at Cibola National Wildlife Refuge for help with sample collection and field logistics.

## Author Contributions

K. Ruegg and T.B. Smith conceived of the study and K. Ruegg wrote the majority of the manuscript and conducted and/or oversaw the analyses. E.C. Anderson wrote the scripts for the population genomic analyses and figure creation. K. Paxton and F. Moore contributed ideas and genetic material for the analysis of migrants from Cibola, AZ. V. Apkenas conducted and helped analyze data for the SNP screening. S. Lao assisted with feather sample organization, extraction, and the analysis of genotyping reliability scores. R.B. Siegel and D.F. DeSante facilitated the collection of feather samples in collaboration with bird banding stations within and outside of the Monitoring Avian Productivity and Survivorship (MAPS) and the Monitoreo de Sobrevivencia Invernal (MoSI) networks.

